# Development of a Transcriptional Biosensor for Hydrogen Sulfide that Functions under Aerobic and Anaerobic Conditions

**DOI:** 10.1101/2025.02.19.639182

**Authors:** Matthew T. Fernez, Shanthi Hegde, Justin A. Hayes, Kathryn O. Hoyt, Rebecca L. Carrier, Benjamin M. Woolston

## Abstract

Hydrogen sulfide (H_2_S) is a gaseous gut metabolite with disputed effects on gastrointestinal health. Monitoring H_2_S concentration in the gut would provide insight into its role in disease, but is complicated by sulfide’s reactivity and volatility. Here we develop a transcriptional sulfide biosensor in *E. coli*. The sensor relies on enzymatic oxidation of sulfide catalyzed by a sulfide:quinone reductase (Sqr) to polysulfides, which bind to the repressor SqrR, triggering unbinding from the promoter and transcription of the reporter. Through promoter engineering and improving soluble SqrR expression, we optimized the system to provide an operational range of 50 µM - 750 µM and dynamic range of 18 aerobically. To enable sensing in anaerobic environments, we identified an Sqr from *Wolinella succinogenes* that uses menaquinone, facilitating reoxidation through the anaerobic electron transport chain by fumarate or nitrate. Use of this homolog resulted in an anaerobic H_2_S response up to 750 µM. This sensor could ultimately enable spatially and temporally resolved measurements of H_2_S in the gastrointestinal tract to elucidate the role of this metabolite in disease, and potentially as a non-invasive diagnostic.

## Introduction

Hydrogen sulfide (H_2_S) is a gaseous metabolite produced by many microbes that inhabit the GI tract^1^. Sulfide has long attracted the attention of researchers due to its unclear, and at times conflicting, role in human GI health and disease^2,3^. Inflammatory Bowel Disease (IBD) affects 1% of the human population with the incidence rate increasing^4^, partially influenced by the widespread popularization of the Western Diet^5^. Beyond genetic predispositions^6^, the onset of IBD is not well characterized mechanistically but it is suspected that microbial dysbiosis may be an agonist^7^. Specifically, metagenomics analysis reveals that sulfate-reducing bacteria (SRB) and enzymes involved in converting cysteine to H_2_S are more prevalent in IBD cohorts than healthy controls^8^. As a molecule genotoxic to enterocytes, H_2_S is suggested to have a deleterious role in inflammation through pro-inflammatory cytokine secretion as a response to DNA damage^9^. Sulfide may also contribute to mucus layer disruption through reduction of disulfide bonds in the mucin network^10^. However, there is contradictory evidence to support a beneficial role in barrier stability at low levels in ulcerative colitis (UC) mice models^11^. Overall, the prevailing emerging view is that the effects of sulfide on host health are highly dose-dependent^12^.

Given the dose-dependence of sulfide’s physiological effects, accurate measurements of its concentration *in vivo* are crucial. However, its reactivity and volatility present major technical barriers, resulting in wide estimate ranges for gut H_2_S levels (25 µM – 2 mM)^13^. Additionally, physiological measurements have traditionally relied on indirect measurements of stool samples and breath. However, these fail to capture spatial variation in sulfide levels that may occur longitudinally throughout the gut or between the lumen and mucosa. There has been extensive research in electrochemical methods to sense sulfide^14–16^, and a miniaturized capsule system with wireless transmitter was recently developed^17^, but their use *in vivo* has not been reported.

Diagnostic bacteria engineered with transcriptional biosensors have recently emerged as an attractive route for non-invasive *in vivo* monitoring of a range of gut metabolites^18^. In these systems, a transcription factor responsive to the target ligand is used to control the expression of a reporter protein. Fluorescent reporters provide convenient readout through analysis of stool samples by flow cytometry^19^. For more stable readout, sensors can be engineered regulate DNA recombinases, encoding the sensing event in the bacteria’s DNA^20^. More recently, coupling a luminescence reporter with a miniaturized luminescence detecting and transmitting electronics in an ingestible micro-bio-electronic device was shown to enable non-invasive real-time readout in pigs^21^. Advances in long-wavelength luminescence reporters also potentially enable the use of biosensors in whole animal imaging studies to probe specific metabolites with spatial and temporal resolution^22–25^. Finally, transcriptional sensors can be used to actuate a therapeutic response, resulting in a “Smart Probiotic” capable of delivering its payload specifically at sites where the level of the target biomarker indicates disease^26^.

Given the benefits of microbial biosensors and the challenges with traditional measurements of H_2_S, we sought to engineer a transcriptional sulfide biosensor. The ideal sensor would meet the following design parameters: 1) An operational range spanning physiological gut concentrations (25 µM - 2 mM); 2) Functionality in both aerobic and anaerobic conditions, owing to the steep radial oxygen gradient from the mucosa to the lumen^27^ and the increased concentration of oxygen in sites of inflammation^28^; 3) A GRAS microbial chassis that could eventually be used diagnostically or therapeutically.

## Results & Discussion

### Sensor Design

Our strategy was to construct an *E. coli-*based H_2_S sensor by adapting the H_2_S-responsive regulatory components previously characterized in *Rhodobacter capsulatus*^29^. This microbe uses sulfide as an electron donor for photosynthesis, and differentially regulates multiple genes in response to exogenous sulfide. The key regulator is the repressor protein SqrR, which binds the *sqr* promoter (P_sqr_). In the presence of sulfide, basal expression of sulfide:quinone oxidoreductase (Sqr) catalyzes the oxidation of sulfide to a persulfide. The formed persufide reacts with two conserved cysteine residues on SqrR to form a tetrasulfide bond that induces a conformation change leading to unbinding and derepression of P_sqr_. Our target host was *E. coli,* given its status as a GRAS microbe and established precedent of using strains such as Nissle 1917 strain in probiotic applications^30^. To adapt the system to *E. coli,* we used two plasmids with complementary origins of replication. (**Figure 1a**). The first plasmid encodes the sensing module (P_tetR_:*sqrR*) and reporting module (P_sqr_:*gfp*) in inverse orientation, separated by a bi-directional terminator. The second plasmid contains the oxidation module, with the *sqr* gene under control of the arabinose promoter (P_bad_). This design allowed us to separately optimize *sqr* expression and the persulfide sensing. Initial tests with the system as depicted yielded no detectable response to H_2_S (Data not shown). We therefore set about optimizing each of the heterologous sensing components for expression in *E. coli*.

**Figure 1.**
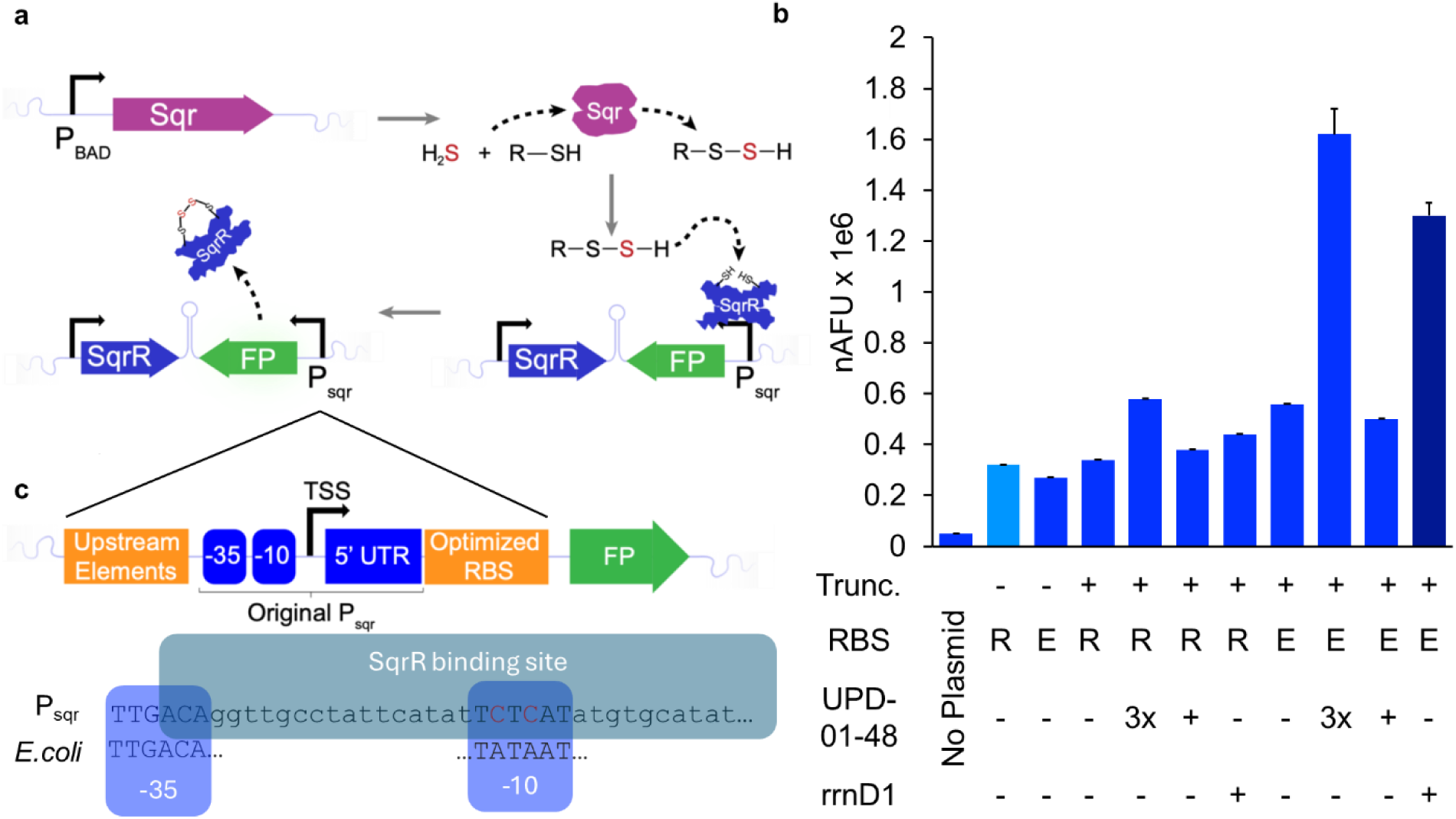
Genetic Architecture of the H2S Biosensor and Engineering to Enhance P_sqr_ Activity. a) H2S is oxidized to polysulfide by a heterologous sulfide:quinone oxidoreductase (Sqr) expressed from a titratable inducible promoter. Exposed cysteines on the repressor SqrR form a tetrasulfide bridge with the formed polysulfide, resulting in dissociation from the P_sqr_ promoter and transcription of the reporter. b) GFP fluorescence from the original and modified P_sqr_ promoters in the absence of SqrR, showing combinatorial effects of truncation (Trunc.), new RBS (E for *E. coli* and R for *R. capsulatus*), and inclusion of upstream elements UP-D01-48 or rrnD1. A serendipitous cloning error resulted in two additional constructs with 3 tandem UP-D01-48 sequences (3x) The original construct is shown in light blue, and the final optimized in dark blue. c) Detailed view of the P_sqr_, showing modifications made to improve activity, including truncation to a core promoter, use of upstream elements and an optimized ribosome binding site (RBS). nAFU = OD-normalized fluorescence. Bars show mean values of n=3 biological replicates with error bars showing the standard deviation.

### P_sqr_ Engineering

We first focused on characterizing and engineering the performance of P_sqr_. Experiments with a control plasmid with GFP under control of P_sqr_ in which the SqrR repressor was removed resulted in minimal GFP expression (**Fig.1b**), suggesting low activity of the native promoter in *E. coli*. One strategy would be to modify the -10 and - 35 boxes to be closer to the consensus *E. coli* sequences. However, previous DNA footprinting experiments revealed that the repressor binding site straddled the -10 box^29^ (**Fig. 1c**), thus mutating this region would risk losing repressor binding. Instead, we tested combinations of three different alterations: First, the originally cloned P_sqr_ sequence included a region of 81 additional base pairs upstream of the -35 box, which we deleted in case it contained unknown regulatory elements. Second, previous work has identified that upstream elements that preceed the -35 box can help recruit Sigma factors to recruit polymerase^31^. We screened two of these, rrnD1 and UPD-01-48^32^, to assess their impact on GFP expression. Lastly, using the ribosome binding side (RBS) RBS calculator^33^, we designed a new sequence to replace the native *R. capsulatus* to improve translation rates. Each of these modifications were screened individually, and in combination. The resultant impact on GFP expression is shown in **Figure 1c**.

Individually, these changes had minor impacts on promoter strength. However, the combination of truncation, new RBS, and UP-D01-48 resulted in a 1.5-fold increase in fluorescence compared to the original construct. Changing the upstream element to rrnD1 with the other two changes outperformed UP-D01-48, yielding a 4.5-fold increase in fluorescence compared to the original promoter. Interestingly, a version of the construct with three tandem repeats of UP-D01-48 derived from a serendipitous assembly product during cloning resulted in a 5.5-fold increase in fluorescence. Given that repeat sequences can be unstable^34^ and that fluorescence from this construct was only marginally brighter than the rrnD1 version, we selected the latter for sensor development.

### Sqr Characterization

With the reporting module in hand, we next confirmed the functionality of Sqr, which catalyzes the oxidation of sulfide into polysulfides for binding with SqrR. Sqr activity in whole cells was quantified using cyanolysis^35^. As shown in **Figure 2a**, the enzyme was highly active, converting 70% of sulfide to polysulfide within 1 hour. We then sought to identify the specific products of the reaction, as Sqr can use a variety of compounds as the sulfane sulfur acceptor, including sulfide, sulfite, and low-molecular-weight thiols like glutathione (GSH)^36^. Given the high concentration of GSH in *E. coli* ^37^, we suspected that GSH was the predominant co-substrate. Cell lysates were derivatized with monobromobimane and analyzed by LC-HRMS, revealing glutathione per-and trisulfide as the only products with an increased peak area compared to empty-vector and no-sulfide controls (**Figure 2b,c** **Supplementary Table 3)**.

**Figure 2.**
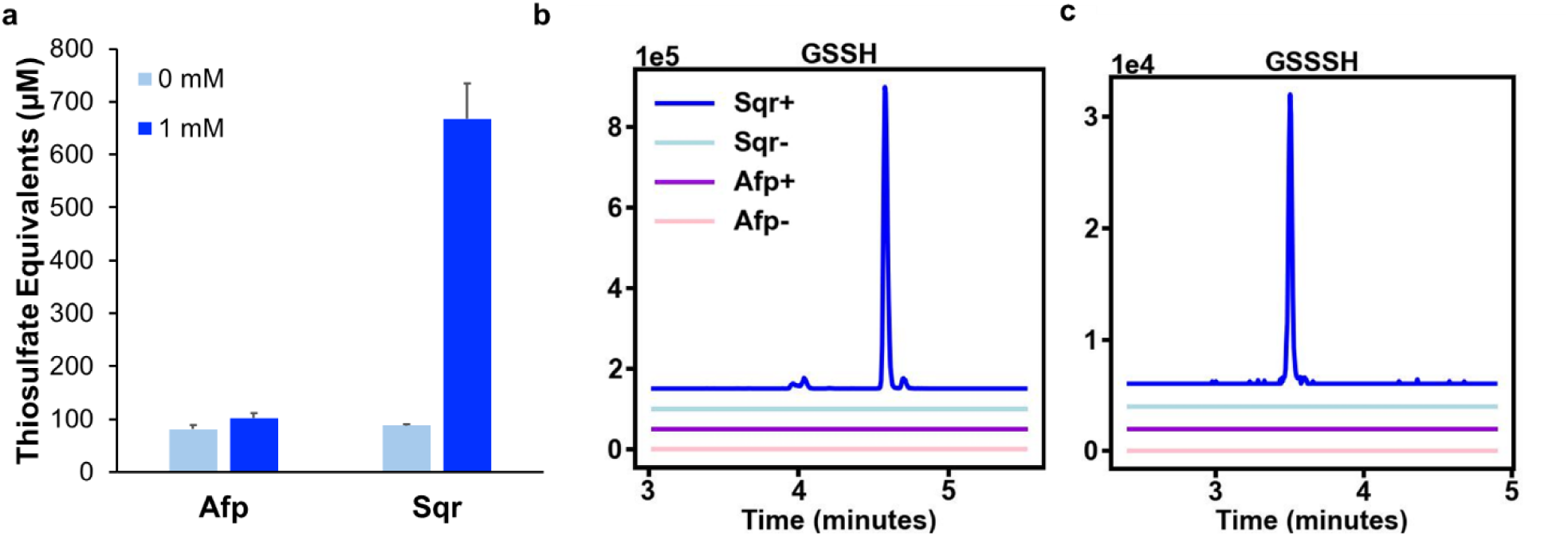
Sqr from *R. capsulatus* produces glutathione persulfide and glutathione trisulfide from Na_2_S. a) Quantification of polysulfides by cyanolysis. Cells harboring pBAD:sqr (Sqr+) or pBAD:mAmetrine (Empty Vector) were grown to OD_600_ 2.0 and resuspended in HEPES, before being treated with 1 mM Na_2_S (bright blue) or vehicle (light blue) and assayed after 1 hour. b and c) LC-HRMS of cell lysates from a) after derivatization with monobromobimane. b: Glutathione persulfide (GSSH), XIC of m/z 530.14. c: Glutathione trisulfide (GSSSH), XIC of m/z 562.09. Data shown are averages of n = 3 biological replicates with error bars showing one standard deviation.

### SqrR Soluble Expression

We next optimized the sensing module. The initial sensor designed used P_Tet_ for inducible expression of SqrR to allow titration of the repressor. Unfortunately, GFP expression from the P_Tet_:SqrR P_Sqr_:GFP expression at full induction was no different than control constructs without SqrR, suggesting low expression of the repressor. We replaced P_Tet_ with P_J23100_, a strong constitutive promoter, but this still resulted in no repression of P_sqr_ promoter activity (**Figure 3**), suggesting the problem might not be the level of expression, but the solubility of SqrR. In previous work, SqrR was purified with an N-terminal His6-SUMO tag to assist in protein folding^29^. We therefore explored constructs containing either His_6_-SUMO, or a His_6_ tag alone, which has also been shown to improve soluble expression^38^. Both tag configurations resulted in soluble expression, as indicated by complete repression of GFP expression in the absence of sulfide (Fig. 3). Excitingly, addition of 750 µM Na_2_S in cells expressing Sqr resulted in an 8.2-fold depression. Since the derepressed expression level was more consistent with the His_6_-SUMO tag, we selected this one for the final sensor design.

**Figure 3.**
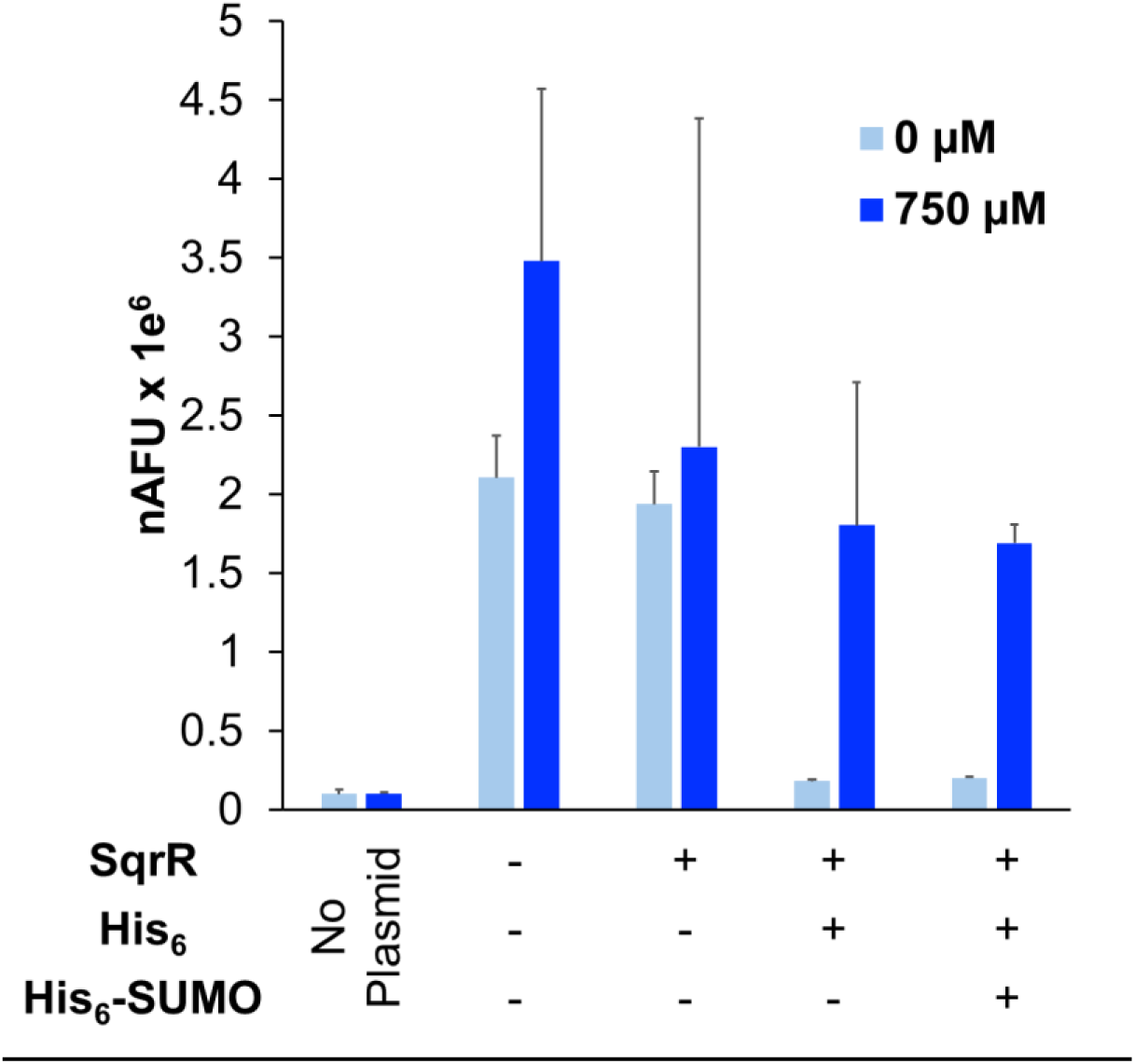
Enhancing SqrR Solubility with N-Terminal Tags Results in Sulfide-Dependent Regulation of P_Sqr_. Bars represent mean OD-normalized fluorescence 16 hours after addition of 750 µM sulfide (bright blue) or vehicle (light blue) in n = 3 biological replicates. Error bars represent one standard deviation.

### Aerobic Sensor Performance

With the individual components optimized, we next characterized the full sensor’s performance. While microtiter plate-based assays are convenient for biosensor optimization, the volatility of sulfide in the slightly acidic culture conditions (pK_a_ = 7.0) resulted in no GFP fluorescence when cells were treated with sulfide in 96-well plates (Data not shown). By contrast, cultures sealed in Hungate tubes with a headspace of air demonstrated a modest sulfide-dependent response up to 250 µM sulfide (**Supplementary** Fig. 1). Sqr couples sulfide oxidation to reduction of a quinone, and sustained activity relies on re-oxidation of the quinol by oxygen through the electron transport chain. We hypothesized that oxygen availability in the Hungate tubes may limit oxidation of higher concentrations of sulfide, reducing the operational range. To test this, we increased the surface area and headspace-to-volume ratio of the cultures by switching from Hungate tubes (5mL culture per 17 mL tubes) to serum bottles (10 mL culture per 120 mL bottle). Excitingly, this increased the upper limit of the operational range from 250 µM to 750 µM (**Supplementary Figure 1**), at a slight cost of dynamic range (6.2 vs. 4.7).

With a working sensor in hand, we investigated additional opportunities to expand the dynamic range. By switching the reporter to mKate, dynamic range increased 35% (**Supplementary Figure 2**), presumably because of decreased background autofluorescence, so we used mKate for all subsequent work. At this point, the sensor only recovered 40% of the hypothetical dynamic range compared to a SqrR-negative control. We hypothesized that the strong promoter (J23100) driving *sqrR* was producing repressor in excess of the intracellular persulfide concentration, even at high levels of H_2_S. To test this, we titrated *sqrR* expression, reducing the promoter strength by approximately 28% and 76% with J23104 and J23105, respectively. Excitingly, the dynamic range increased from 8 to 18 with the J23105 promoter (**Figure 4a**). Most intriguingly, the fluorescence values of the J23105-based sensor above 250 µM sulfide are higher than in the unrepressed control lacking sqrR (**Figure 4a**). Both the original J23100-based sensor and the J23105 variant exhibited a sigmoidal dependence on sulfide, with K_A_ of 91 ± 16 µM and n of 2.7 ± 1.3 for J23100, and K_A_ of 193 ± 12 µM and n of 2.5 ± 0.3 for J23105.

**Figure 4.**
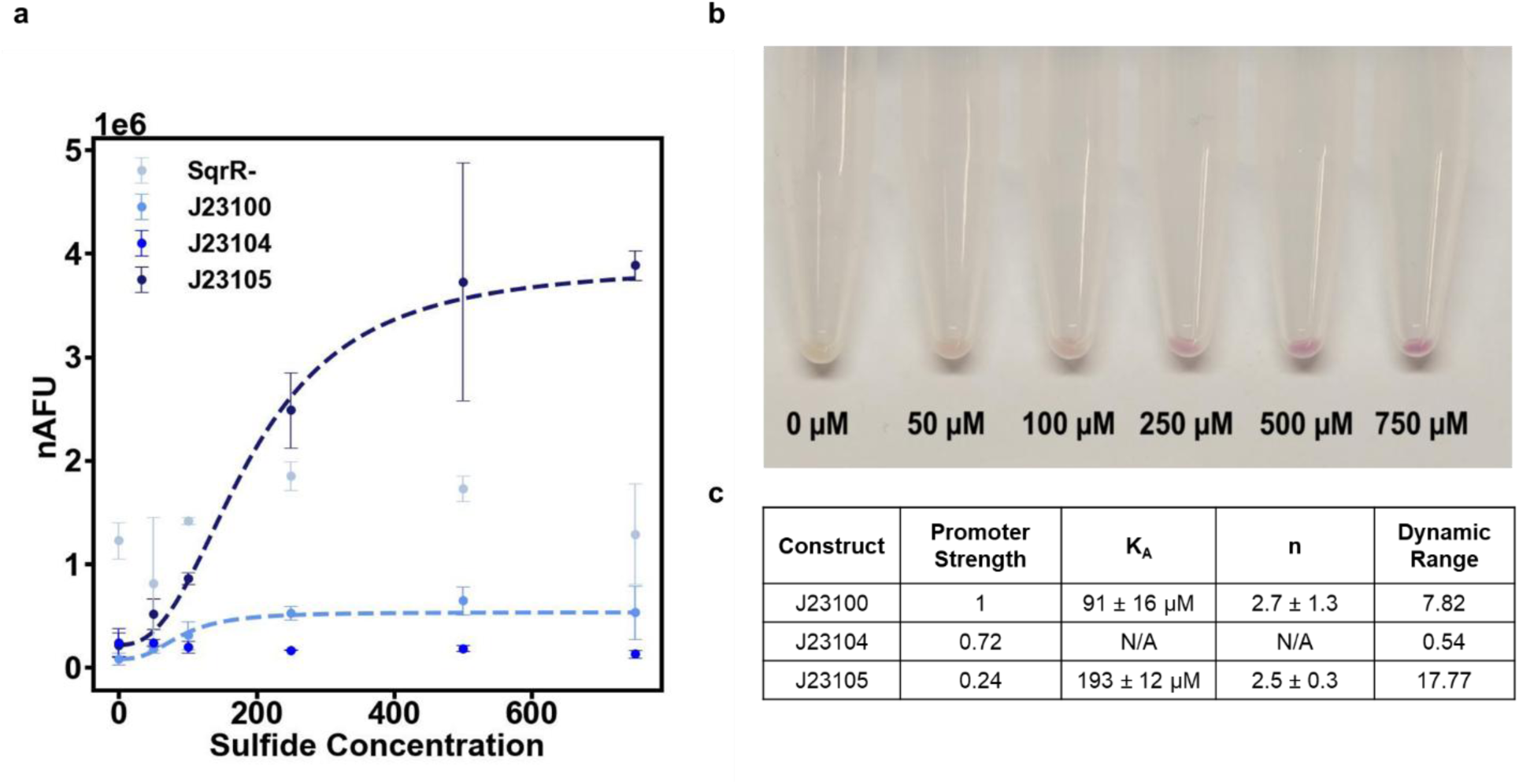
SqrR titration improves sensor performance. (a). OD-normalized fluorescence 16 hours after sulfide addition for various sensor constructs as a function of concentration. For J23100 and J23105, Hill parameters were fitted to the experimental data and curves shown as dotted lines. Data points are means of n = 3 biological replicates. Error bars are +/- one standard deviation. (b). Pellets from the J23105-based sensor with sulfide exposure increasing left to right, showing visible increase in red color from mKate. (c). Table of parameters for the 3 sensor constructs, showing half-maximal ligand occupation (KA), cooperativity (n) and dynamic range. Hill parameters are presented with 95% confidence intervals.

### Engineering the Sensor to Work Anaerobically

As described above, sustained Sqr activity depends on continuous reoxidation of the quinol pool through the electron transport chain. In the previous experiments, oxygen served as the terminal electron acceptor, but the gut microbiota is primarily anaerobic. Fumarate and nitrate are two alternative electron acceptors found in the GI tract, with fumarate abundant in the healthy gut^39^, and elevated nitrate associated with inflammation^40^. Both are reduced in *E. coli* by quinol-dependent reductases. We thus hypothesized that the sulfide sensor should function anaerobically in the presence of these compounds. Unfortunately, in contrast to previous observations^41^, in our hands the *Rhodobacter* Sqr showed minimal activity anaerobically with either of these electron acceptors by cyanolysis assay (Fig. 5a).

**Figure 5.**
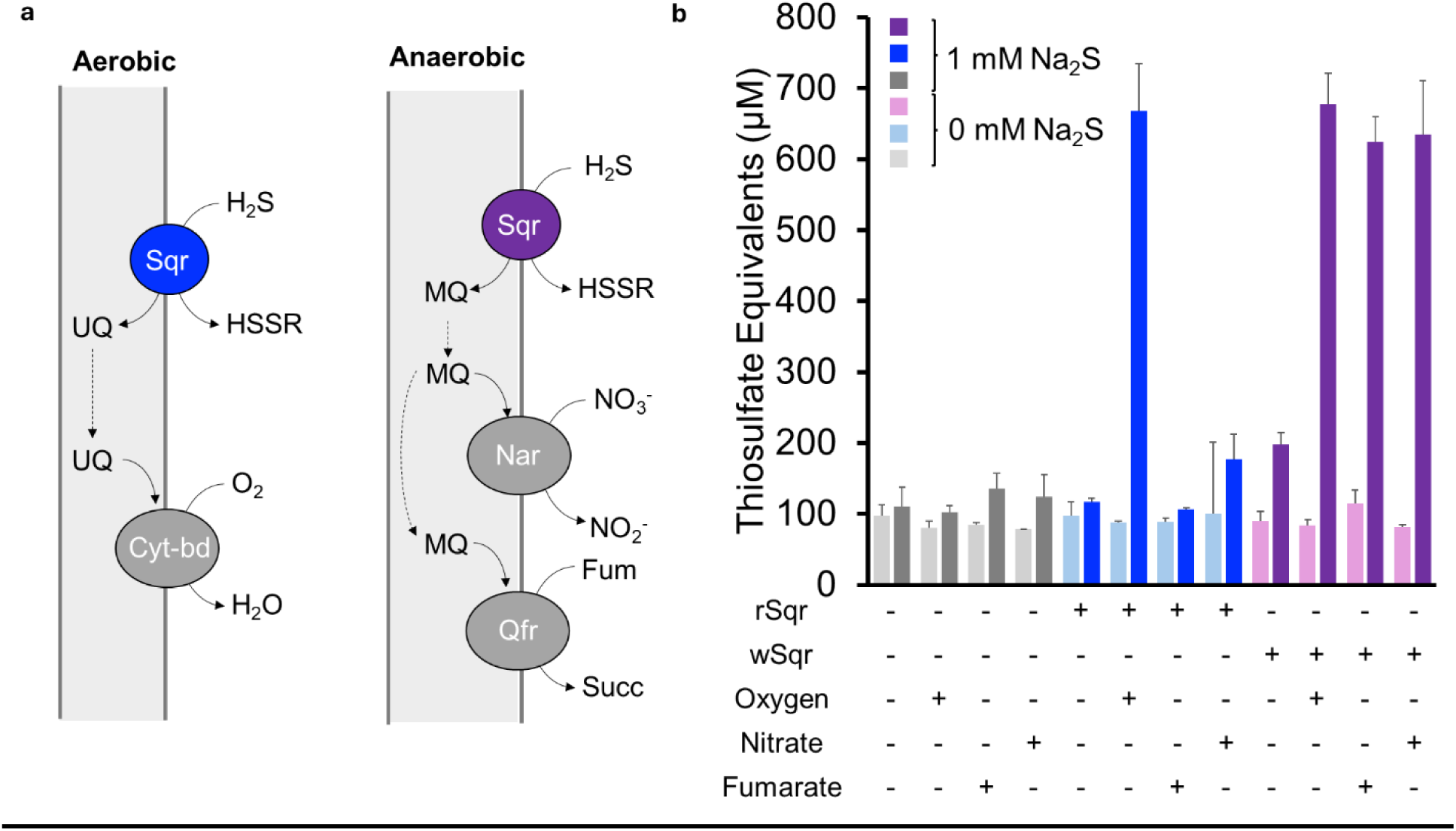
Sqr from *Wolinella succinogenes* enables anaerobic conversion of sulfide to polysulfides. a) Schematic of electron transfer from sulfide to terminal different electron acceptors until aerobic (left) or anaerobic (right) conditions. b) Quantification of Sqr activity by cyanolysis with the *Wolinella* (wSqr) and *Rhodobacter* (rSqr) homologs under aerobic conditions and in anoxic environments with or without 20 mM sodium nitrate or sodium fumarate. Bars represent means of n = 3 biological replicates, with error bars representing one standard deviation. Abbervations: UQ, ubiquinone; MQ, menaquinone; HSSR, polysulfide; Cyt-bd, cytochrome oxidase; Nar, nitrate reductase; Qfr, quinone:fumarate reductase

We suspected this may be due to a mismatch between the quinone specificity of Sqr and the quinones available during aerobic and anaerobic growth (Fig. 5b). Under aerobic conditions, ubiquinone is the dominant quinone in *E. coli*^42^, but under anaerobic conditions menaquinone becomes the primary^43^. We hypothesized that the *Rhodobacter* Sqr may have minimal activity with menaquinone, preventing robust anaerobic activity, consistent with the role of this enzyme in aerobic sulfide oxidation^44^. To overcome this challenge, we sought to identify an alternative Sqr that used menaquinone as a redox cofactor. Given that every characterized fumarate reductase is menaquinone-dependent ^45^, we reasoned that an Sqr from an organism that could couple sulfide oxidation to nitrate reduction would likely be menaquinone-specific.

*Wolinella succinogenes* is an obligate anaerobe that was shown to couple sulfide oxidation to fumarate reduction for growth^46^, though an Sqr was neither identified nor characterized. Using the *Rhodobacter* Sqr (rSqr) sequence as a BLAST query, we identified an Sqr homolog in *W. succinogenes* (Accession # WP_011138184.1). The corresponding gene was amplified from *Wolinella* gDNA and the putative Sqr was tested under both aerobic and anaerobic conditions with fumarate and nitrate available as anaerobic electron acceptors. Remarkably, the *Wolinella* Sqr (wSqr) could perform sulfide oxidation with a similar yield under both aerobic and anaerobic conditions (Fig. 5b).

The capability of *Wolinella* Sqr (wSqr) to oxidize sulfide in *E. coli* under anaerobic conditions was promising for developing an anaerobic biosensor. We next incorporated this gene into the oxidation module of the sensing strains, and tested the sensor anaerobically, using aerobic fluorescence recovery to detect mKate fluorescence^47^. As shown in **Figure 6**, the anaerobic sensor with the wSqr responded over a similar range of sulfide concentrations, with slightly different dynamics. With fumarate in the culture media, the sensor produced a binary response: While there was no increase in fluorescence between 50 and 250 µM (**Supplementary Figure 3**), there was a statistically significant increase observed at concentrations above 500 µM, with a similar dynamic range as the aerobic sensor recovered (15.7 vs. 18, respectively). With nitrate, there was an overall lower dynamic range (5.7), but there appeared to be a more graded but statistically insignificant concentration-dependent response (**Supplementary Figure 3**).

**Figure 6.**
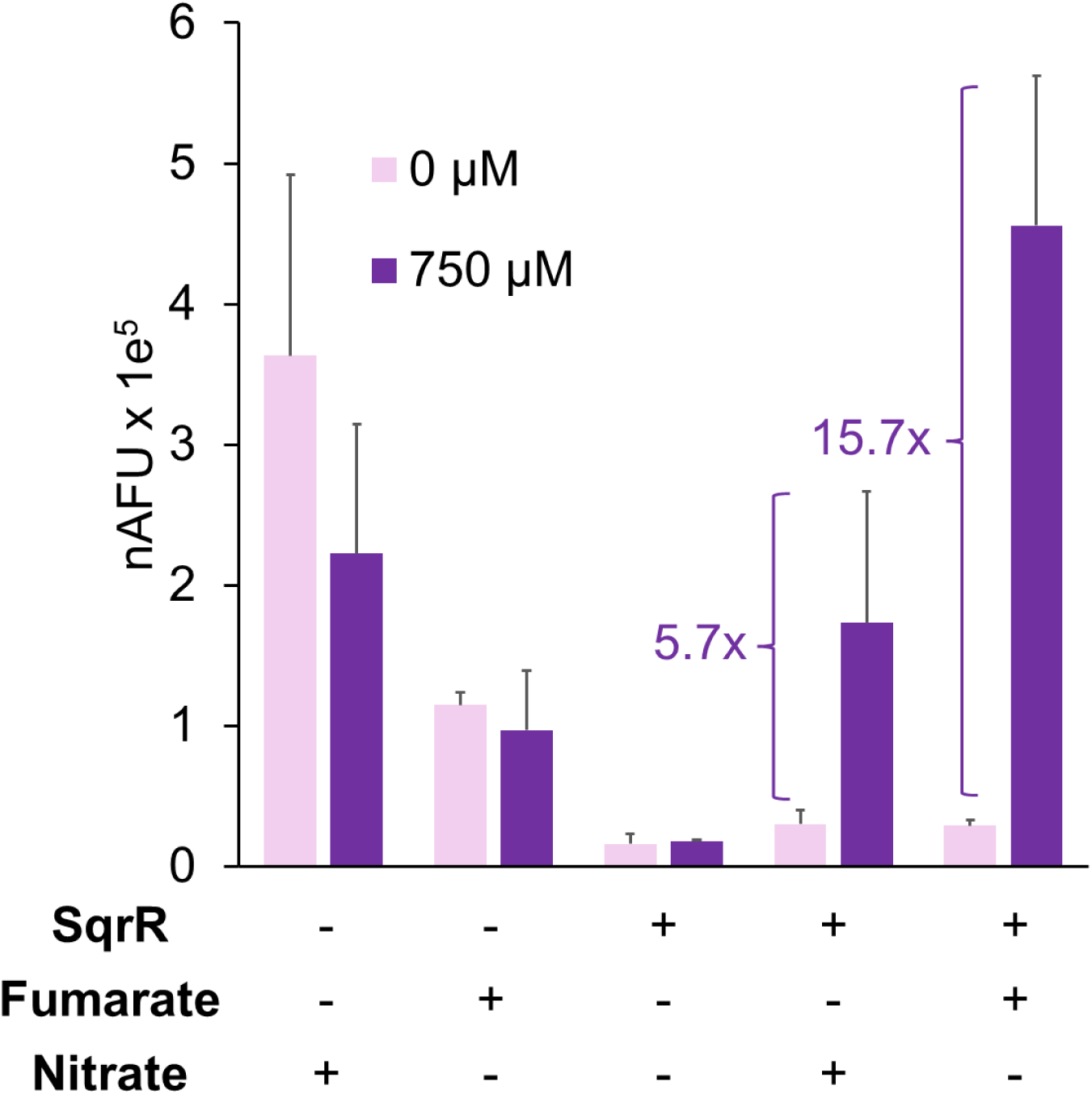
Anaerobic sensor discriminates between physiologically high and low levels of sulfide. Normalized fluorescence values of the wSqr-based sensor under anaerobic conditions with either fumarate or nitrate, and with or without 750 µM sodium sulfide. Bars represent means of n=3 biological replicates with error bars of one standard deviation.

The dynamic range of the sensor with supplemented fumarate was 15.7, on par with the output using rSqr in aerobic conditions. Interestingly, as with the aerobic sensor, the fully induced expression is higher than the no-SqrR control. The reason for the limited resolution of intermediate sulfide concentrations is presently unclear. However, the sensor can clearly delineate between physiologically low and high levels, which could eventually be useful in a clinical or diagnostic context, especially given the challenges with current methodologies for H_2_S detection.

To the best of our knowledge, this work is the first report of a transcriptional biosensor for detection of exogenous sulfide under both aerobic and anaerobic conditions. A previous report^48^ detailed construction of a similar reporter consisting of the persulfide-responsive CstR repressor and cognate promoter from *Staphylococcus aureus,* and Sqr from *Cupriavidus pinatubonensis* or *Pseudomonas putida.* However, fluorescence output from the reporter strain saturated at 10 µM H_2_S, which is in the range produced by WT *E. coli.* Thus, the authors focused instead on using their sensor as an elegant quorum-sensing system. As shown in Figs. 5 and 6, our sensor is responsive to much higher concentrations of sulfide, allowing it to reliably detect different levels of exogenously supplied sulfide within the physiologically relevant range found within the gut^13^. We speculate that the key difference is the relative sensitivity of *CstR* and *SqrR* to persulfide, given these proteins are from different families^29^.

Given its ability to detect elevated levels of sulfide both aerobically and anaerobically, the sensor developed here may have promise as a non-invasive diagnostic tool. Additionally, our demonstration of a strain capable of anaerobically oxidizing sulfide could eventually lead to new therapeutics based on engineered probiotics, given that compromised sulfide oxidation capacity is associated with IBD^49^. The sensor could also find use in other anoxic environments where sulfide plays an important physiological role, such as the tumor microenvironment^50^. In the future, the sensor could be engineered into probiotic strains designed to modulate H2S levels in vivo^51^.

Beyond the practical utility of the sensor, an additional interesting finding from this work was the importance of the HIS_6_-SUMO tag on the functionality of the repressor. Metagenomic screening has emerged as a promising method of identifying new transcription factors responsive to a wide range of chemical ligands^52^. Such screens should consider the use of such tags to enhance solubility of heterologous regulators to avoid false negatives. The use of UP elements to enhance transcription from a suboptimal promoter may also prove a general strategy for engineering novel transcriptional machinery into heterologous hosts. Overall, this work provides a sensitive transcriptional sulfide biosensor that could be used in diagnostic situations, as well as novel approaches to the optimization of biosensors based on heterologous transcription factors.

## Materials and Methods

### Media and Chemicals

All chemicals were purchased from Fisher Scientific unless otherwise stated. Bacterial strains were cultured in either Luria-Bertani (LB) medium containing 5 g/L yeast extract, 10 g/L tryptone, and 10 g/L NaCl or a supplemented M9 medium (M9+). M9+ contained 6 g/L Na_2_HPO_4_, 0.5 g/L NaCl, 3 g/L KH_2_PO_4_, 1 g/L NH_4_Cl, 1 mM MgSO_4_, 0.1 mM CaCl_2_, 4 g/L glucose, 1 g/L Casamino acids and 1 mg/L Thiamine-HCl. Carbenicillin (100 µg/L) and spectinomycin (60 µg/L) were used for plasmid maintenance, and chloramphenicol used for growth inhibition during aerobic fluorescence recovery experiments at 20 µg/L. Induction of *sqr* was achieved through addition of 10 mM L-arabinose.

### Plasmid Construction and Strains

*E.coli* s1030 was chosen as the final sensor strain, as it carries a genomic copy of *tetR* to enable anhydrotetracycline (aTc) induction, and constitutive *araE* expression for titratable arabinose induction^53^. All cloning was performed in *E.coli* DH5a (New England Biolabs, NEB). Medium copy-number plasmids pTR47 (SpR) and pTR48 (AmpR)^54, 55^ were chosen for the sensor cassette and Sqr expression, respectively. PCR and Gibson assembly were done using Q5 Polymerase and Hi-Fi Assembly Mix (NEB). Constructs were verified by sequencing (Azenta/Primordium). DNA and plasmid purification kits were purchased from Zymo Research. *Rhodobacter capsulatus* genes were codon harmonized^56^ and synthesized as gBlocks by Integrated DNA Technologies. Promoter configurations were sampled through overlap extension PCR from the Andersen library series (iGem Part:BBa_J23100) for *sqrR* expression. HIS_6_-SUMO tag, GFP, mKate, pBAD elements were PCR-amplified from the plasmids shown in Table S1. *Wolinella succinogenes* was purchased from DSMZ and the *sqr* gene was amplified from genomic DNA. Final plasmids were transformed into chemically competent *E. coli* s1030 using heat shock^57^. Single colony isolates were selected off antibiotic agar plates and grown in LB overnight before cryopreserved in 20% glycerol at -80 ⁰C. All plasmids and primer sequences can be found in **Table S1** and **Table S2** respectively. The final biosensors constructs are available from AddGene

### Sensor Assay

#### Aerobic Sensor Experiments

Overnight cultures were diluted 1:100 in fresh M9+ medium in shake flasks at 200 RPM with carbenicillin, spectinomycin, and L-arabinose. Cells were grown at 37⁰C until OD 0.3, then 10 mL were aliquoted into 120 mL serum bottles. Sodium sulfide (Na_2_S) - prepared as a stock solution at 100 mM in 100 mM NaOH – was added to the desired concentration, and cultures quickly sealed with butyl rubber septa and aluminum crimp seals to minimize H_2_S evaporation. Sulfide assays were conducted at this point to confirm the initial concentration. Vessels were then returned to the shaker for 12-18 hours, after which 200 µL samples were transferred to a 96-well plate for measurements of OD_600_ and fluorescence (SpectraMax i3, Molecular Devices). GFP and mKate measurements were taken at wavelength pairings 485/515 and 585/635 respectively.

#### Anaerobic Sensor Experiments

For anaerobic experiments, pre-cultures were grown anaerobically from cryostocks in M9+ medium in Hungate tubes overnight. Following the same dilution as above, serum bottles were prepared in the anaerobic chamber using degassed medium and electron acceptors (20 mM sodium nitrate or 20 mM sodium fumarate) and transferred to the shaker. After cells reach OD_600_ 0.3, they are returned to the chamber and split into 6 individual serum bottles (10mL) for each sulfide level following the same timeline as stated above and received sulfide from stock solution prepared anaerobically. For fluorescent sampling, GFP matured in minutes following withdrawal by syringe and needle, and were measured immediately. Samples taken from mKate matured more slowly, and were treated with chloramphenicol to inhibit protein synthesis during a 2-hour aerobic maturation prior to recording the final fluorescence value.

All sensor outputs are reported as normalized Arbitrary Fluorescence Units (nAFU), which is media blank-adjusted fluorescence output divided by media blank-adjusted OD_600_:

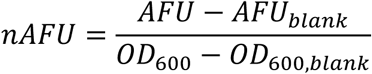

Where appropriate, sensor data was fit to the below Hill equation using scipy curve_fit in Python for plotting and parameter estimation.

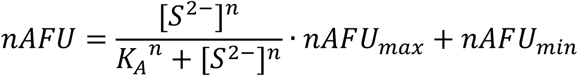

Where *nAFU_max_* is the fluorescence at the highest level of sulfide, and *nAFU_min_* is fluorescence with no sulfide.

### Methylene Blue Sulfide Assay

To confirm sulfide concentrations in liquid samples, we used the previously adapted methylene blue assay modified for 96-well plate format^51^. Briefly, 200 µL of cell culture sample was added to a mixture containing 600 µL of 1% (w/v) zinc acetate and 15 µL of 3 N NaOH, and vortexed. Following a 5-minute incubation period, 150 µL of 0.1% n-dimethylethlenediamine in 5N HCl and 150 µL of 23 mM ferric chloride in 1N HCl were added. Samples were then centrifuged at 16,000 g for 5 minutes. 200 µL of supernatant were transferred to a 96-well plate for absorbance measurement at 670 nm, and compared to a standard curve prepared in the identical medium.

### Hot Cyanolysis for Polysulfide Quantification

To quantify Sqr activity, a cyanolysis procedure was adapted from^35^. Overnight cultures were diluted 1:100 in LB and grown to OD 0.3 before adding 10 mM L-arabinose and cultured for an additional 3 hours. Cells were harvested by centrifugation at 3,000 g for 10 minutes and washed in 1/10 culture volume of 50 mM HEPES (pH 7.0) before resuspension to OD_600_ of 2.0 in the same buffer. 3 mL of cell suspension was then aliquoted into Hungate tubes, treated with 1 mM Na_2_S and shaken at 37⁰C for 1 hour before sample collection. 250 µL of cell suspension was added to pre-prepared microcentrifuge tubes containing 550 µL of 1% boric acid (w/v in water) and 200 µL of 100 mM potassium cyanide. Samples were then boiled using a heat block at 100 ⁰C for 5 minutes, and cooled to room temperature. Then, 100µL of ferric nitrate reagent (3 g of ferric nitrate in 5 mL of 33% perchloric acid) was added to each sample and centrifuged at 16,000 g for 5 minutes. 200 µL supernatants of samples were transferred to a 96-well plate and absorbance was recorded at 460 nm. Quantification was achieved through comparison to a standard curve of sodium thiosulfate, processed the same way but with the addition a of 5 µL of 1 M copper sulfate as a catalyst for thiosulfate conversion.

### Identification of Sqr Products by LC-HRMS

1.5 mL of cells from the cyanolysis experiments above were centrifuged and resulting pellets were frozen at -80⁰C for analysis of reaction products by LC-HRMS following derivatization with monobromobimane (mBBr) ^58^. Lysis and derivatization was accomplished in a single step using a lysis buffer consisting of 50% v/v acetonitrile in water with 20 mM ammonium bicarbonate and 5 mM mBBr, prepared fresh daily and covered in foil to prevent light exposure. Cell pellets were resuspended in 200 µL of mBBr lysis buffer and mixed by pipetting. Samples were incubated at 60 ⁰C for 1 hour in the dark with a heat block covered in foil. Samples were then centrifuged at 20,000 g for 10 minutes to precipitate cellular debris. Supernatants were 5-fold diluted in LCMS grade water, then filtered via syringe (0.2 µm filter) into vials. Samples were analyzed on an Agilent 6545 LC-MS QTOF equipped with an electrospray ionization (ESI) source, and an Agilent Zorbax Eclipse Plus C18 UPLC column. Mobile phase A was 0.1% formic acid in water, mobile phase B was 95% acetonitrile with 5% water, and all solvents were LCMS grade. The gradient started at 98% mobile phase A, and ramped to 2% mobile phase B over 7.5 minutes, at 0.4 mL/minute with a column temperature of 40 ⁰C. The QTOF was operated in positive mode, using a fragmentor voltage of 125 V. The MS scan range was 100 m/z to 1700 m/z, with a resolution of approximately 20,000 for 500 m/z. Predicted compound formulas of polysulfide products were used with the Find By Formula tool in Agilent’s MassHunter software to identify polysulfide peaks. Find By Formula identified the peaks using the exact mass, natural isotope spacing and abundance predicted from the compound formula. Retention times were obtained from derivatized standards where possible.

## Author Contributions

MTF, BMW, and RLC conceived this project. MTF and BMW designed the constructs and experiments. MTF carried out the experiments and analyzed the data. SH helped with cloning and testing of *Wolinella* Sqr constructs. JAH optimized and assisted in chemical sulfide measurements. MTF and BMW prepared and revised the manuscript. All authors approve the final version of the manuscript.

## Acknowledgements

The authors thank David Liu’s lab for donation of pTR47 and pTR48. This work was also advanced by the contributions from undergraduates Brielle Quigley, Ella Sweet, and Julia Chen, who assisted in sensor experiment execution. The authors are grateful for financial support to BMW from an NIH NBIB Trailblazer (R21EB033892) and to RLC from an NIH NIBIB R01 (1R01EB021908). Figures were prepared with BioRender.

## Abbreviations

IBD: Inflammatory Bowel Disease
GI: Gastrointestinal
H_2_S: Hydrogen Sulfide
Sqr: sulfide:quinone (oxio)reductase
SRB: sulfate-reducing bacteria
UC: ulcerative colitis
nAFU: normalized arbitrary fluorescence units
LC-HRMS: Liquid chromatography-high resolution mass spectrometry
mBBr: monobromobimane,
TSS: transcription start site
RBS: ribosome binding site.

