## Supplementary Material for "Development of a Transcriptional Biosensor for Hydrogen Sulfide that Functions under Aerobic and Anaerobic Conditions"


1. Supplementary Table 1- Plasmids and Strains
2. Supplementary Table Note 1- DNA Sequences in this Study
3. Supplementary Table 2- Oligos for PCR/Assembly
4. Supplementary Table 3- Sqr Products via LCMS
5. Supplementary Figure 1- Hungate Tubes vs Serum Bottles
6. Supplementary Figure 2- GFP vs mKate
7. Supplementary Figure 3- Expanded Anaerobic Sensor Data

**Supplementary Table 1- Plasmids and Strains**

| Name | Description | Reference |
|---|---|---|
| *Strains* | | |
| *E. coli* DH5α | Cloning strain | NEB |
| *E. coli* S1030 | Sensor host. Expresses *araE* and *tetR* for $P_{BAD}$ and $P_{Tet}$ induction by L-arabinose and anhydrous tetracycline, respectively | Liu lab[1] |
| *Wolinella Succinogenes* | Source of putative *sqr* homolog amplified and expressed in PMF2 for anaerobic sulfide oxidation | DSMZ 1740[2] |
| *Plasmids* | | |
| pMF1 | Original sensor architecture. pSC101 origin of replication. $P_{Tet}$-SqrR and $P_{sqr}$-GFP. *SqrR* codon harmonized. *ampR* for ampicillin family resistance | This study, [3] |
| pMF1.R | Replacement of SqrR with RFP for $P_{sqr}$ optimization | This study |
| pMF1.1R | Truncation of extraneous base pairs in $P_{sqr}$ before -35 region | This study |
| pMF1.2R | RBS attached to $P_{sqr}$ -GFP replaced with thermodynamically calculated RBS | This study |
| pMF1.3R | Truncation from pMF1.1R and RBS from pMF1.2R | This study |
| pMF1.1R_1r | Insertion of rrnD1 element before -35 box in pMF1.1R | This study |
| pMF1.1R_2u | Insertion of UPD148 element before -35 box in pMF1.1R | This study |
| pMF1.1R_2u3 | Insertion of 3 UPD148 element (tandem repeat) in pMF1.1R (cloning oddity) | This study |
| pMF1.3R_1r | Insertion of rrnD1 element before -35 box in pMF1.3R. Chosen as final $P_{sqr}$ permutation | This study |

| | | |
|---|---|---|
| pMF1.3R_2u | Insertion of UPD148 element before -35 box in pMF1.3R | This study |
| pMF1.3R_2u3 | Insertion of 3 UPD148 element (tandem repeat) in pMF1.3R. Yielded highest fluorescence but not chosen over repeat sequence concerns | This study |
| pMF1.3_1r | Replacement of RFP with SqrR in pMF1.3R_1r | This study |
| pMF1.3_1r_J1 | Replacement of $P_{Tet}$ with J23100, constitutive promoter form Anderson library | This study, iGem Part BBa_J23100 |
| pMF1.3_1r_J1H | Insertion of N-terminal $HIS_6$ tag on SqrR | This study |
| pMF1.3_1r_J1HS | Insertion of N-terminal $HIS_6$-SUMO tag on SqrR. Best repression phenotype, tag configuration chosen | This study |
| pMF1.3R_1r_J1 | Replacement of SqrR with RFP in pMF1.3_1r_J1 for $P_{sqr}$GFP positive control strain | This study |
| pMF1.3_1r_J1HS_mK | Replacement of GFP with mKate as fluorescent reporter of sulfide | This study |
| pMF1.3G_1r_J1_mK | Replacement of SqrR with GFP and GFP with mKate in pMF1.3R_1r_J1 as the positive control (SqrR-) for mKate sensors. | This study |
| pMF1.3_1r_J4HS_mK | Replacement of J23100 in pMF1.3_1r_J1HS_mK with J23104 to reduce promoter strength | This study, iGem Part:BBa_J23104 |
| pMF1.3_1r_J5HS_mK | Replacement of J23100 in pMF1.3_1r_J1HS_mK with J23104 to reduce promoter strength | This study, iGem Part:BBa_J23105 |
| pMF2-rSQR | Enzymatic plasmid, $P_{BAD}$ driven *sqr* codon harmonized from *R. capsulatus.* Contains aaDa | This study, [3] |

| | | |
|---|---|---|
| | for Spectinomycin resistance and colE1 origin of replication | |
| pMF2-wSQR | Native *sqr* from *W. succinogenes* replaces *Rhodobacter sqr* | This study, [2] |
| pMF2-AFP | Control strain for *sqr* plasmids. mAmetrine (AFP) replaces *sqr*. | This study |

**Supplementary Note 1: Sequences included in this study:**

Native $P_{sqr}$ Sequence from *R. capsulatus*:

CCTGACGCAAATTCATGCTAAGGTTGTGGGGCGGACGCTGGTCCGCCCCTTTCTT
TTTCGCCTGATCCGCGGGACCGATCTTCCCGCTTGACAGGTTGCCTATTCATATTC
TCATATGTGCATATACGGGCTTCGGCCCGACAGCCAGTTCGGGGAGGGACAG

Codon-Harmonized *sqr* from *R. capsulatus*:

ATGGACACAGCCCAAGATCCGCAAGATGACTTTGACCCGGAGATGGGGTCAGATA
CGGATGAACGCTGCGCGGCGCTGGATGCCGAAGAGATGGCGACACGTGCGCGTG
CGGCCAGCAACCTGCTTAAAGCGCTGGCGCACGAGGGCCGTCTAATGATTATGTG
CTATCTTGCCAGCGGCGAAAAGAGCGTGACCGAACTGGAGACCCGTCTGTCAACG
CGTCAGGCGGCGGTTTCGCAGCAACTGGCGCGTCTGCGTCTTGAAGGGCTGGTG
CAATCGCGACGCGAAGGCAAAACGATTTATTATAGCCTTTCAGACCCGCGCGCCG
CCCGCGTGGTGCAGACGGTTTACGAACAGTTTTGCAGCGGCGATTAA

Final, engineered $P_{sqr}$:

CCGCGGTAGAATTCCAGAAAAAAAGATCAAAAAAATACTTGACAGGTTGCCTATTC
ATATTCTCATATGTGCATATACGGGCTTCGGCGACCTGTAAAAATTATAAATTTTTTT
AAGGAGGTTTTTT

$HIS_6$-SUMO Tag:

ATGAAATCTTCTCACCATCACCATCACCATGGTTCTTCTATGGCTAGCATGTCGGAC
TCAGAAGTCAATCAAGAAGCTAAGCCAGAGGTCAAGCCAGAAGTCAAGCCTGAGA
CTCACATCAATTTAAAGGTGTCCGATGGATCTTCAGAGATCTTCTTCAAGATCAAAA
AGACCACTCCTTTAAGAAGGCTGATGGAAGCGTTCGCTAAAAGACAGGGTAAGGA
AATGGACTCCTTAAGATTCTTGTACGACGGTATTAGAATTCAAGCTGATCAGACCC
CTGAAGATTTGGACATGGAGGATAACGATATTATTGAGGCTCACAGAGAACAGATT
GGTGGG

Synthesized mAmetrine (AFP) gBlock:
ATGGTTTCTAAAGGGGAGGAACTTTTTACAGGGGTCGTTCCTATTCTTGTAGAGTT
GGATGGGGACGTAAACGGCCACAAATTCTCGGTGCGTGGTGAGGGTGAGGGAGA
CGCCACAAATGGGAAGCTTACCCTTAAATTCATTTGTACCTCAGGGAAATTGCCTG
TCCCATGGCCCACGTTGGTGACTACCCTGAGTTACGGCGTACAATGCTTCGCACG
CTATCCGGACCACATGAAGCAACACGATTTCTTCAAGTCTGCGATGCCGGAGGGG
TATGTCCAGGAGCGTACAATCTCGTTTAAGGATGATGGGTCTTATCGCACTCGCGC
AGAGGTTAAATTTGAGGGCGACACGCTGGTAAATCGCATCGAGTTAAAAGGCATC
GATTTTAAGGAGGATGGTAACATCCTTGGGCACAAGTTAGAATACAACATGAATGT
GTGGGACGCATATATCACAGCCGATAAACAAAAGAATGGAATCAAGGCCAATTTCA
AAATTGAGCATAACGTGGAGGATGGCGGAGTCCAGTTAGCCGACGCGTATCAGCA

AAACACGCCCATCGGGGACGGGTCGGTTTTACTGCCCGATAATCACTATTTATCAT
TCCAGTCTAAATTATTCAAGGACCCGAACGAACAGCGTGACCATATGGTATTACTG
GAGTTCGTGACCGCAGCCGGGATTACACCTGGTATGGATGAATTGTATAAG

Native *Wolinella sqr*:

ATGTCCTCACTTGATAGGGAATGGCTAGAGGCTTTAGAGCAGATGGATGGGGAGC
TCAAAAAGGCGGGGTTGAGCCGAAGGGATGCGCTCAAGGTTCTAGGGCTTGGAG
GGGCGGCTCTCGCTCTCCCTGCGACCGCTCCCAGGGCGCACGCGGCGAGTAATG
CTAAAGGGAAGATTGTCATTATTGGAGCAGGACTCGCAGGAATCACTGTGGCATC
GCGTCTAAGCCACGCCCTTAGCAAGCCTGATATCACGATCATTGATGGGGGCGAT
AAGGTCGATTACCAACCTGGATACACATTGATTGCCTCGGGTGTTTATGGGCCCAA
TGATGTCACTTACGAGCGTGCAGGACTCATTCCTAGTGGTGCGAAGTGGATCAAA
GAGTATGTCAAAGAGATTGATGCGGCAGGAAATAGCGTCACCACCACCTCAGGTC
AAAAGATCACTTATGACTATCTCGTGGTTGCGACAGGTTTGGTGACTGATTATTCAA
TGGTTAAAGGTTTGAGCGAAAGCGACGTGGGTCGCAATGGAATCGCCTCTATCTAT
ACGCTTCCCACCTGCCAAAAGGCCTTTGGCCAAATCAAGGAGTTTGTCAACCAGG
GGGGTGTAGGGCTTTTCACCGACCCCCATACTCCTATTAAATGCGGAGGTGCTCC
CAAAAAGATTCAGTTTCTCGTGGATGACTACGCCCGAAAAGAGGGGAAACATGACA
AGATAAAGACGATCTTTCTGCCCAATGGGGGGACAATGTTTGGGGTCAAAGAGTA
CGCAGAGATGATTGAACGCCTCTACAAAGAGAAGGGGATGGAGTGGAAATTCAAG
CACAATCTCGTGGCGATCGATCCCGCTTCCAAAAAGGCGACCTTTGAGTACACCTA
CACAGTCAAGGGAGAGTTTGATGAGGTGTTGGGAGAGCATGAGCTCATTACCCAA
AAAGATAATGTGGTGATGGATTATGACTTTATCCATGTCACCCCTCCGATGAGAGC
CCCCAAGATGGTTAAAGATTCGGAGCTCTCTTGGAAGAGGGGAAGTGCATCAGCT
GGGGGCTGGATGGAGCTTGTGAAGGAGACCTTGCAACACCCTGTCTATAAAAATG
TCTTTGGTCTTGGTGATGTGGCGGGAATCCCCATGGGTAAAACAGGGGGAAGCGT
AAGGAAGCAAGCGCCTGTTTTGGTGGAGAATCTCATTAACGCGATGGAAGGCAAA
GAGCCCACCGCTCAATATGGCGGCTACACCGTATGCCCTCTGATTGTCGATTATG
GTCGTGTGGCGATGCTAGAATTTGACTGGAGCGCCACACCCAAGCCCTCCTTCCC
TCTTGATCCTTCAGTGCCTAGGTGGGTTTATTGGGCGATGAAAGTCTATATGCTGA
AACCTATGACCATGATTGGGATGCTTAAAGGCTACGCCTAA

**Supplementary Table 2: Oligos in this study**

| Name | Sequence | Description |
|---|---|---|
| oMTF_001 | GGATCTTGGGCTGTGTCCATTTTTTTTTCCTCCTTATAAGGGG | Amplification of pTR47 backbone for pMF1 |
| oMTF_002 | TAGCATGAATTTGCGTCAGGCCCCCCTATAGTATATAACACTG | Amplification of pTR47 backbone for pMF1 |
| oMTF_003 | GCCAGTTCGGGGAGGGACAGATGAGCAAGGGCGAAGAGCT | Amplification of GFP for pMF1 assembly |
| oMTF_004 | AGTTTTGCAGCGGCGATTAAACAGTCAGTAGGGCCCTAAA | Amplification of GFP for pMF1 assembly |
| oMTF_005 | AAGCCGTTAAGAAAGGATAGACTTAATTAACGGCACTCCT | Amplification of pTR48 backbone for pMF2 assembly |
| oMTF_006 | AGTACCACAATATGAGCCATTTTTTTTTCCTCCTTAGCTC | Amplification of pTR48 backbone for pMF2 assembly |
| oMTF_016 | TTTAGGGCCCTACTGACTGTTTAAGCACCGGTGGAGTG | Amplification of RFP for pMF1.R |
| oMTF_017 | CTTATAAGGAGGAAAAAAAAATGGCGAGTAGCGAAGAC | Amplification of RFP for pMF1.R |
| oMTF_018 | TTTTTTTTCCTCCTTATAAGGGGG | Amplification of pMF1 backbone for pMF1.R |
| oMTF_019 | ACAGTCAGTAGGGCCCTAAAAAAAAAC | Amplification of pMF1 backbone for pMF1.R |
| oMTF_020 | GTGCATATACGGGCTTCGGCGACCTGTAAAAATTATAAATTTTTTTAAGGAGGTTTTTTATGAGCAAGGGCGAAGAGC | Insertion of RBS |
| oMTF_021 | GCCGAAGCCCGTATATGCACATATG | Insertion of RBS |
| oMTF_024 | TATAGGGGGGCCGCGGGACCGATCTTCCCGCTTG | $P_{sqr}$ truncation |
| oMTF_025 | AAGATCGGTCCCGCGGCCCCCCTATAGTATATAACACTGG | $P_{sqr}$ truncation |
| oMTF_026 | GTGTTATATACTATAGGGGGGCCGCGGTAGAATTCCAGAAAAAAAGATCAAAAAAATACTTGACAGGTTGCCTATTC | rrnD1 insertion |
| oMTF_027 | GTGTTATATACTATAGGGGGGCCGCGGGGAAAATTTTTTTTAAAAAAATCGTGCTTGACAGGTTGCCTATTC | UPD148 insertion |

| | | |
|---|---|---|
| oMTF_028 | CCGCGGCCCCCCTATAGTATATAACAC | Reverse primer for either upstream element insertion |
| oMTF_061 | TGATGGTGAGAAGATTTCATTTTTTTTTCCTCCTTATAAGGGGGTAG | Insertion of $HIS_6$ tag |
| oMTF_062 | ACAGAGAACAGATTGGTGGGATGGACACAGCCCAAGATCCGCA | Insertion of $HIS_6$ tag |
| oMTF_063 | TGCGGATCTTGGGCTGTGTCCATCCCACCAATCTGTTCTCTGT | Insertion of $HIS_6$-SUMO tag |
| oMTF_064 | CTTATAAGGAGGAAAAAAAAATGAAATCTTCTCACCATCACC | Insertion of $HIS_6$-SUMO tag |
| oMTF_081 | CTAGTATTTCTCCTCTTTCACTAGTAGCTAGCACTGTACCTAGGACTGAGCTAGCCGTCAATCATCTGGCCATTCGATGGAC | Replacement of $P_{Tet}$ with J23100 |
| oMTF_082 | ACTAGTGAAAGAGGAGAAATACTAGATGGACACAGCCCAAGATCCGC | Replacement of $P_{Tet}$ with J23100 |
| oMTF_084 | ACTAGTGAAAGAGGAGAAATACTAGATGAAATCTTCTCACCATCACC | Replacement of $P_{Tet}$ with J23100 for $HIS_6$-SUMO SqrR |
| oMTF_085 | ACTAGTGAAAGAGGAGAAATACTAGATGGCGAGTAGCGAAGACGTTATC | Replacement of $P_{Tet}$ with J23100 for RFP |
| oMTF_105 | TTGGGCTGTGTCCATAGAAGAACCATGGTGATGGT | Insertion of $HIS_6$ tag on J23100-sqrR |
| oMTF_105 | CACCATGGTTCTTCTATGGACACAGCCCAAGATC | Insertion of $HIS_6$ tag on J23100-sqrR |
| oMTF_113 | CTAGTATTTCTCCTCTTTCACTAGTAGCTAGCACAATACCTAGGACTGAGCTAGCTGTCAATCATCTGGCCATTCGATGGAC | Replacement of J23100 with J23104 |
| oMTF_114 | CTAGTATTTCTCCTCTTTCACTAGTAGCTAGCATAGTACCTAGGACTGAGCTAGCCGTAAATCATCTGGCCATTCGATGGAC | Replacement of J23100 with J23114 |
| oMTF_138 | ACGGAGCCAATGTACGC | Replacement of GFP with mKate |
| oMTF_139 | AAAAAACCTCCTTAAAAAAATTTATAATTTTTACAGGTC | Replacement of GFP with mKate |
| oMTF_143 | TTTGCGTACATTGGCTCCGTTTATCTGTGCCCCAGTTTGC | Amplification of mKate for sensor |
| oMTF_144 | TTTTTTTAAGGAGGTTTTTTATGTCTGAGCTGATTAAGGAG | Amplification of mKate for sensor |
| oMTF_147 | GCTCTTCGCCCTTGCTCATCCCACCAATCTGTTCTCTGT | Replacement of RFP with GFP for mKate positive control strain |
| oMTF_148 | GGATGAACTCTATAAGTAAACAGTCAGTAGGGCCCTAAAAAAAAAC | Replacement of RFP with GFP for mKate positive control strain |

| | | |
|---|---|---|
| oMTF_149 | GGGCCCTACTGACTGTTTACTTATAGAGTTCATCCATGCC | Replacement of RFP with GFP for mKate positive control strain |
| oMTF_150 | CACAGAGAACAGATTGGTGGGATGAGCAAGGGCGAAGAG | Replacement of RFP with GFP for mKate positive control strain |
| oMTF_088 | ACTTAATTAACGGCACTC | Amplification of pMF2 backbone for AFP or other *sqr* genes |
| oMTF_089 | TTTTTTTTCCTCCTTAGCTC | Amplification of pMF2 backbone for AFP or other *sqr* genes |
| oSH_001 | GAGCTAAGGAGGAAAAAAAAATGTCCTCACTTGATAGGGAATG | Amplification of Native *Wolinella sqr* |
| oSH_002 | GATGCTTAAAGGCTACGCCTAAACTTAATTAACGGCACTCCT | Amplification of Native *Wolinella sqr* |

**Supplementary Table S3. Sqr Product Candidates and LCMS Hits**

| Compound | Formula | Predicted m/z | Retention Time (min) | Peak Area: Sqr Sulfide + (m/z found) | Peak Area: Sqr Sulfide - | Peak Area: Afp Sulfide + | Peak Area: Afp Sulfide - |
|---|---|---|---|---|---|---|---|
| Hydrogen Sulfide Bimane Adduct | $C_{20}H_{23}N_4O_4S$ | 415.1440 | 3.60 | 3883165 (415.1458) | 90175 (415.1449) | 197148 (415.1453) | 42162 (415.1456) |
| Cysteine Bimane Adduct | $C_{13}H_{18}N_3O_4S$ | 312.1018 | 1.99 | 1646 (312.0886) | 5136 (312.0909) | 3550 (312.0885) | 2321 (312.0904) |
| Glutathione (GSH) Bimane Adduct | $C_{20}H_{28}N_5O_8S$ | 498.1658 | 2.6 | 487535 (498.1611) | 74905 (498.1611) | 32580 (498.1614) | 47485 (498.1616) |
| Glutathione Persulfide (GSSH) Bimane Adduct | $C_{20}H_{28}N_5O_8S_2$ | 530.1379 | 4.57 | 1951828 (530.1500) | *0** | *0** | *0** |
| Glutathione Trisulfide (GSSSH) Bimane Adduct | $C_{20}H_{28}N_5O_8S_3$ | 562.1100 | 3.5 | 63490 (562.0943) | *0** | *0** | *0** |

***No hits at specified retention time***

### Supplementary Figure 1- Serum bottles improves sensor operational range

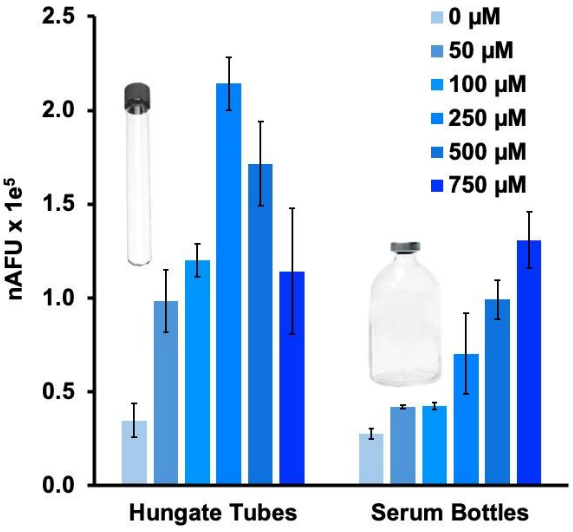


Normalized fluorescence for the J23100-GFP sensor grown in LB taken 16 hours after sulfide addition compared between using Hungate tubes and serum bottles. Data are averages of n=2 with error bars representing +/- one standard deviation.

**Supplementary Figure 2-** Dynamic range increases with use of mKate as the reporter protein

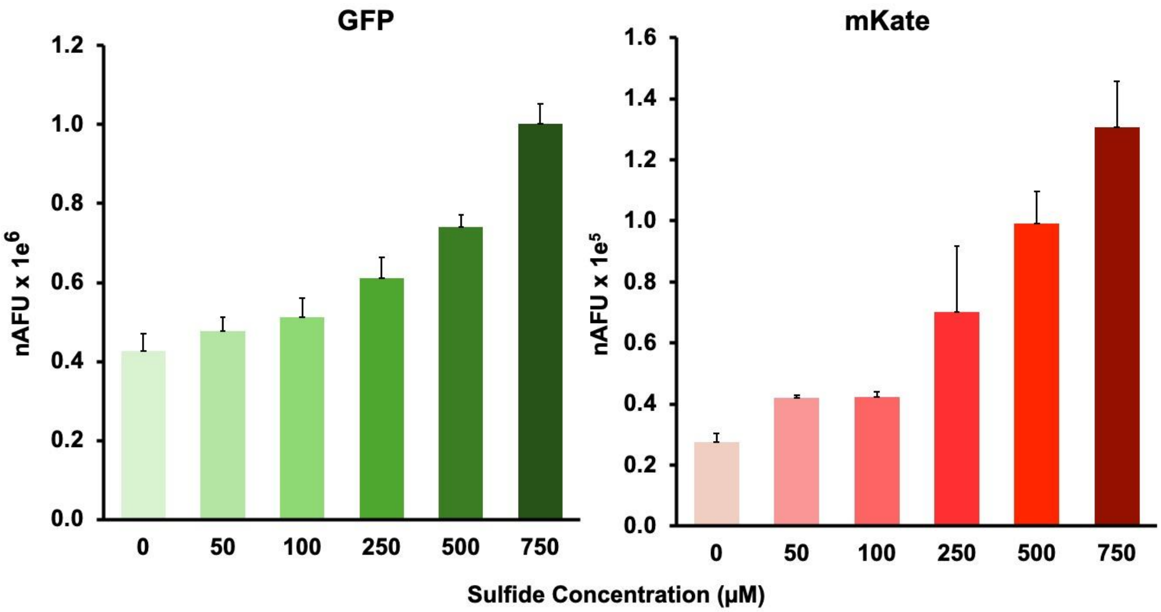


Supplementary Figure 2. Normalized fluorescence of the J23100 sensor with either GFP or mKate as the reporter protein grown in LB in serum bottles taken 16 hours after sulfide addition as a function of sulfide concentration. Data are averages of n=2 with error bars representing one standard deviation above the mean.

**Supplementary Figure 3-** Full range of sensor response under anaerobic conditions

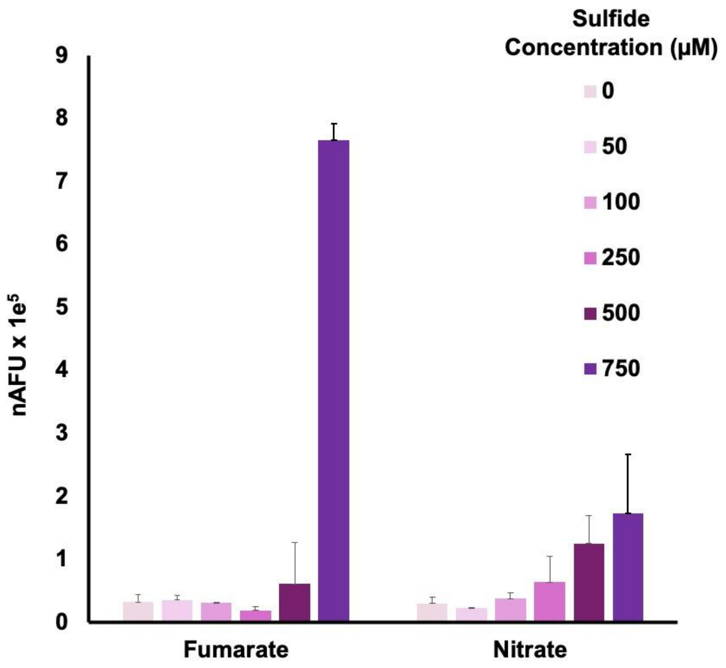


**Supplementary Figure 3.** Normalized mKate fluorescence of anaerobically grown J23105 sensor taken 16 hours after sulfide addition with either fumarate or nitrate available as a terminal electron acceptor for Sqr. Data are averages of n=3 with error bars representing one standard deviation above the mean.
